# Biomarkers for capturing disease pathology as molecular process hyperstructure

**DOI:** 10.1101/573402

**Authors:** Arno Lukas, Andreas Heinzel, Bernd Mayer

## Abstract

**Background:** Precision of drugs in clinical development but also of approved treatment sees limits, documented as attrition in clinical stage drug testing and suboptimal number needed to treat in clinical practice. Precision medicine aims at approaching a causal relation of disease pathology, treatment mechanism of action and clinical outcome. The instance linking pathology, clinical phenotype and drug response is disease characteristics amenable for quantitation, including established clinical phenotyping parameters and upcoming molecular profiling and biomarkers. Molecular biomarkers situated at the interface of pathology-specific molecular process architecture and drug mechanism of action promise capturing aspects allowing assessment of treatment response.

**Results:** Approximating a set of 1,008 disease terms as pathology molecular networks provides 3,860 molecular processes involving 4,602 protein coding genes. Assembling this process set in a hierarchical cluster using mean shortest paths among processes as distance measure allows representation of molecular processes in cumulative aggregation. This procedure transforms human disease pathology into a static instance of a molecular process hyperstructure involving 1,340 aggregate levels in a molecular architecture. The hyperstructure allows evaluating molecular biomarker candidates at different levels of molecular process aggregation in terms of biomarker-specific entropies. Interpretation as information content reflects the capacity of a biomarker for sensing molecular process configuration.

Deriving entropies across aggregation levels for a reference set of 1,502 biomarker candidates identifies significant spread in information content of individual biomarkers. Exemplified on biomarker panels holding evidence for prognostic capacity and factors serving as drug targets from selected chronic diseases, biomarker entropies allow interpretation in terms of sensitivity for capturing process context and specificity for informing on the status of individual processes afflicted with a given pathology.

**Conclusions:** High entropy biomarkers provide candidate molecular proxies for clinical phenotyping parameters, and low entropy biomarkers add information on specifics of disease pathology. Combining high and low entropy biomarkers in panels may offer relevant resolution of molecular process configurations for improving patient stratification with respect to minimizing variance in drug response.

## Background

Approaching precision in management of diseases has been the central challenge in clinical practice as well as in development of new treatment modalities. However, despite significant advancements in experimental capabilities joined with data rich analytics, the number of new molecular entities (NMEs) approved by the United States Food and Drug Administration (U.S. FDA) remained constant in the last ten years, with in the mean 30 NMEs approved annually [1]. Specifically for age-associated, chronic diseases as certain immune-mediated, metabolic or neurodegenerative disorders, causative treatment is rare, at the same time seeing major attrition in clinical stage drug development programs [2]. Aside challenges in developing novel intervention, post-approval analysis of various drugs demonstrates lack of precision, as expressed by suboptimal number needed to treat (NNT) [3,4].

Evidence-based treatment guidelines establish the gold standard in clinical care, reflecting a quantitative relation of clinical presentation, treatment decision and outcome. Treatment effect size initially grounds in clinical trial design, using inclusion criteria from established clinical phenotyping. Such parameters serve for patient stratification, in case of a successful trial constituting drug label and prescription information. Such stratification is deemed to respect the effect of treatment mechanism of action in terms of benefit in outcome as well as risk for triggering adverse drug reactions.

Clinical practice executes further stratification for adding precision on a personalized level. Such optimization involves for instance adjustment of drug dosing in relation to individual disease factors, or even off-label use of drugs in case of lacking response to standard regimens [5]. Also in certain clinical trial designs a drug response component is explicitly factored in as enrichment element for optimizing stratification via use of surrogate parameters [6]. One such approach evaluated short-term response to a novel medication utilizing surrogate parameters associated with outcome, and only in case of meeting certain response criteria randomizing into treatment and placebo [7]. With recent focus on the term precision medicine the quest is in replacing stratification procedures derived on a cohort level with causation on a patient level. Goal is minimizing variance in drug response while at the same time optimizing benefit. This approach needs tools for decision support, ideally already at initiation of treatment.

Translational and preclinical research provides detailed insight about a drug’s direct molecular target context. Still, specifics of a given patient’s pathophysiology embedding such target remain as central factor driving variation in drug response. While for certain patients a drug target context may be decisive for developing a clinical endpoint, the same context may be less relevant for other patients assigned to the same clinical phenotype according to clinical parameters used. Ideally, phenotyping allows differentiating such pathology specifics. In case of high variability in drug response given clinical phenotyping parameters apparently fall short in defining homogeneous cohorts, in consequence impacting precision in both, clinical trials as well as clinical practice.

Molecular phenotyping has been proposed as promising route for adding accuracy in stratification, covering from genetic profiling to the various omics tracks [8], with molecular biomarkers as central component [9]. However, aside selected examples from clinical trials for patient pathology stratification [10-12] and drug pharmacology aspects [13], molecular phenotyping has hardly made impact so far in clinical routine. The number of regulatory approvals of companion diagnostics aimed at improving stratification even falls short compared to approval of novel medication. Next to optimizing characterization of disease pathologies and identification of relevant drug targets and respective compounds, improved patient phenotyping is of equal relevance for improving precision.

Aside a plethora of proposed biomarker candidates from discovery studies lacking subsequent validation, relevant approaches in most part fostering multi-parameter classification have entered clinical evaluation. Examples cover high dimensional molecular classifiers [10,11] as well as clinical parameter integration efforts [14], together with models involving both clinical and molecular parameters [15]. Most of these advancements focus on improving prognosis on a cohort level. Improved prognosis is of relevance in clinical trial design regarding frequency of accepted study endpoints and in consequence number of patients to be included for meeting statistical power. Pace of disease progression is also relevant in clinical practice when it comes to initiation and specifics of treatment regimens. However, even when stratifying for progression with improved accuracy the question remains if the parameters utilized meet with increased probability for beneficial response to a certain treatment. One perspective follows the assumption of a continuum progression for a given phenotype on a disease population level. Such approach assumes a homogeneous natural course of disease, and selecting fast progressors allows defining a stratum with limited pathophysiological variability.

Equivalently valid and factually becoming evident as variation in drug response in various diseases, a cohort homogeneous in progression may be heterogeneous in underlying pathology. In such case homogeneity is falsely introduced by the phenotype parameters describing progression, although precision regarding treatment effect needs to leverage on accurate response prediction [16]. With these considerations a specific class of phenotype parameters is to be identified, allowing personalized prediction of drug response on the background of disease prognosis.

The following approach evaluates the propensity of molecular biomarker candidates to educate on systemic consequence of disturbance in individual, disease-specific molecular processes. Indication for improving molecular phenotyping is given via combining biomarkers monitoring specific disease pathology aspects together with parameters reflecting the consequence of such specifics on a patient’s molecular pathophysiology level. Such strategy in molecular biomarker panel design may offer optimized stratification with respect to minimizing variance in drug response.

## Results

### Approximating a human pathology hyperstructure

With acknowledging relevance of molecular context for evaluating prognostic and predictive biomarker candidates, molecular network models capturing disease pathologies are considered as valid representation. The National Center for Biotechnology Information Medical Subject Headings (NCBI MeSH) controlled vocabulary of diseases holds in total 4,860 individual disease terms. 3,066 diseases are unique leaf terms according to this vocabulary, reflecting most specific disease phenotypes in the sense of clinical classification. Assigning protein coding genes reported as involved in individual pathologies according to NCBI PubMed identifies in the mean 36 molecular features for a given leaf term. Mapping the molecular feature set of each leaf term on a protein coding gene interactome results in 1,412 disease networks (i.e. 1,654 leaf terms cannot be represented on such graph level due to lack of sufficient assignment of protein coding genes according to the data source used). In a disease molecular network each node codes for a disease-associated molecular feature and the edges indicate molecular interactions. Further segmenting each disease-specific network into clusters of densely connected nodes (k-cores) and demanding for each leaf term presence of at least one k-core holding at least three protein coding genes provides k-cores for 1,008 leaf terms. This procedure approximates pathologies as molecular network modules, covering about 30% of leaf disease terms listed in NCBI MeSH. The average number of k-cores per disease term is 4, and each k-core involves in the mean 10 protein coding genes. The 1,008 disease pathologies are represented by in total 3,860 k-cores, each k-core in the following being considered as molecular processes. 4,602 unique protein coding genes are involved in at least one of the 3,860 k-cores.

We assume that each individual process holds biomarkers which can be used as process status proxy, i.e. serving as observable reporting on characteristics of level 1, **O^1^**, (molecular process level, where in contrast level 0 observables, **O^0^**, reflect characteristics of individual molecular features in isolation without taking into account their specific molecular process involvement). Further, in pathologies each process is to be considered in the context of other processes. Following the notion of pathologies being process-of-processes configurations, a process distance matrix can be derived reflecting the pair-wise shortest path between all processes. Using the underlying protein coding gene interactome as reference network the mean path length between all nodes of one process and node sets embedded in the other processes can be derived. This distance measure provides means for process aggregation in a complete linkage hierarchical clustering, resulting in a static approximation of a molecular process hyperstructure (Fig. 1a).

**Fig 1:**
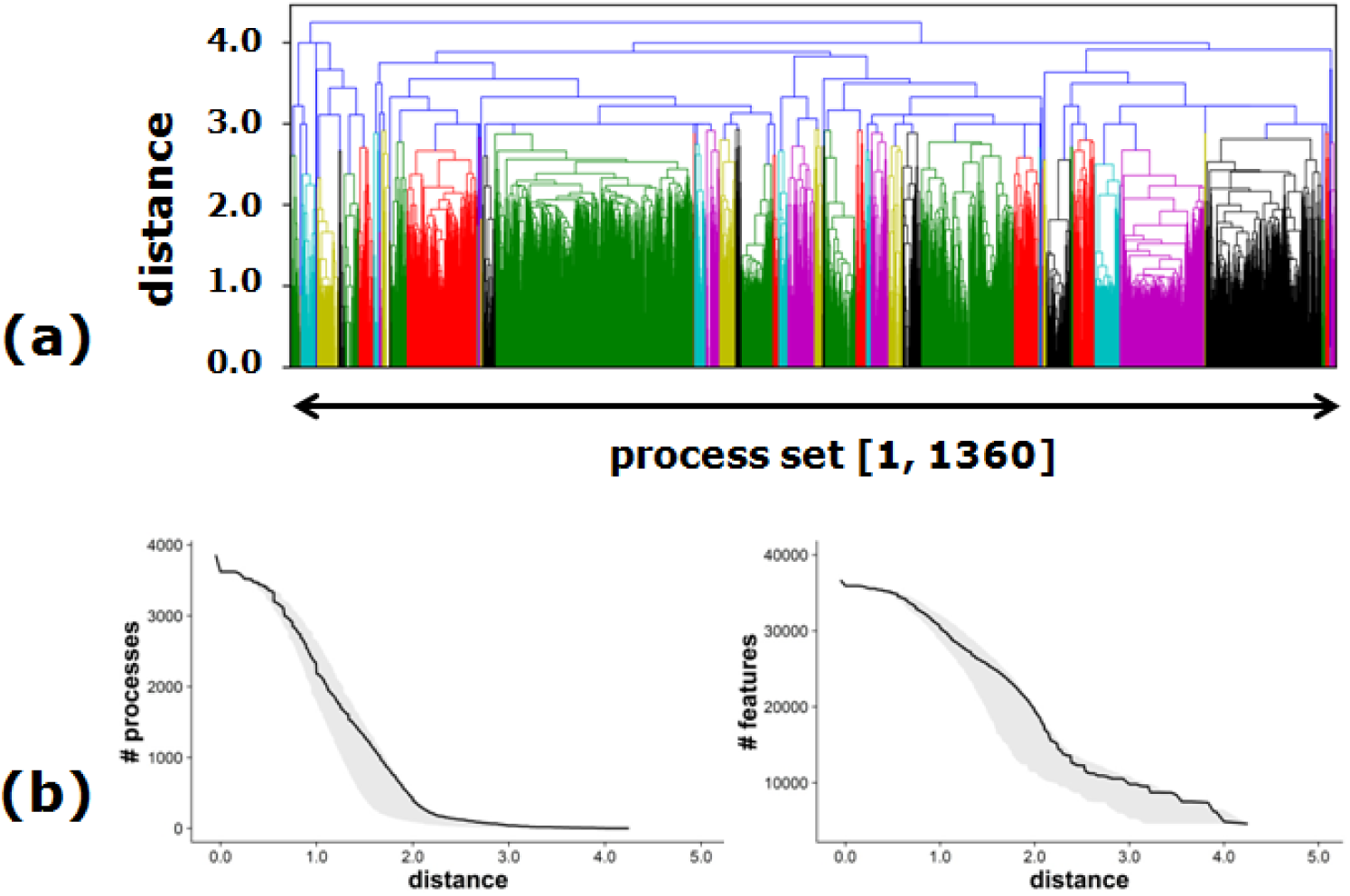
Molecular process aggregation. (a) Dendrogram of process aggregation. The x-axis holds the set of 3,860 molecular processes at level 1, their aggregation being determined as function of mean shortest path distance between processes (y-axis). Color coding indicates clusters of local aggregation. (b) Number of processes (# processes) and number of protein coding genes (# features) according to process aggregation as function of mean shortest path distance. The black line reflects process aggregation referring to the given protein coding gene interactome. The grey area shows aggregation characteristics resting on reference interactomes with reduced connectivity.

Aggregation as given in Fig. 1a displays gradual formation of core process aggregates, at higher distance further merging and incorporating smaller aggregates for finally reaching complete linkage. Both, number of processes as well as number of features approximate a sigmoid shape in dependence of distance (Fig. 1b). As control, interactomes with reduced connectivity via randomly removing interactions are used for evaluating stability of aggregation characteristics. Aside systematic shifts in distance as introduced from diminished connectivity aspects of aggregation remain stable.

With the start set of 3,860 processes (level 1, L1) the first aggregation at a distance zero involves 238 processes. According to given disease molecular annotation this set of processes resembles identical k-cores as process set members of the 1,008 disease terms. Subsequent aggregation constitutes at fast pace up to a distance of 2.1 (aggregation step 1,177), leaving 290 process aggregates to further merge. The highest level in the dendrogram is reached after 1,340 aggregation steps (including equidistant merges at various mean shortest path distances). All processes culminate in a single aggregate (a process-of-processes) holding all 4,602 unique features at a mean shortest path distance of about 4.2. With this procedure a static process hyperstructure across 1,008 human pathologies of order 1,340 is established.

While the set of protein coding genes (level 0) involves 4,602 features, significant redundancy is introduced at L1 (the set of 3,860 processes), embodying 36,682 features in a redundant manner. Feature count at level 1,177 is 17,465, clearly indicating slower pace in decline of feature redundancy when compared to pace of process aggregation.

The hierarchical process configuration can be used for evaluating the information content of biomarkers with respect to serving as proxy of process aggregation levels.

### Information content of biomarkers

Extracting annotation for the descriptors “biological marker, diagnostic/prognostic” from NCBI MeSH for the set of 1,008 disease terms identifies biomarker candidates assigned to 1,520 protein coding genes, with the constraint of each biomarker being member of at least one of the 3,860 processes. In this molecular feature set evidence for serving as biomarker covers from genomic alterations to changes in protein concentration, in the following all assigned to the respective protein coding gene. In total there are 7,012 biomarker-to-disease term assignments (diagnostic and/or prognostic) embedded in the given set of pathologies. The feature with most frequent biomarker annotation is TP53 involved in 210 processes and assigned to 50 disease terms, followed by AKT1 (160 processes, 13 disease terms) and EGFR (147 processes, 30 disease terms). With given constraints on evaluating biomarker coverage specifically in k-cores the disease term holding most biomarker assignments is non-small cell lung carcinoma holding 254 biomarkers. In contrast, 494 of the 1,008 disease terms do not hold any biomarker assignment on the level of k-cores.

According to definition a biomarker serves as proxy for the status of a process, where each individual process itself is embedded in process context as approximated in Fig. 1a. Single assignment of a biomarker at level 1 is interpreted as proxy for a specific process. In contrast, multiple assignment of a biomarker candidate at level L1 consolidates to unique presence only in aggregate processes. Number of aggregation steps needed for seeing a unique assignment of a biomarker to a single aggregate process depends on relatedness of process context according to mean shortest path distance (Fig. 2a).

**Fig 2:**
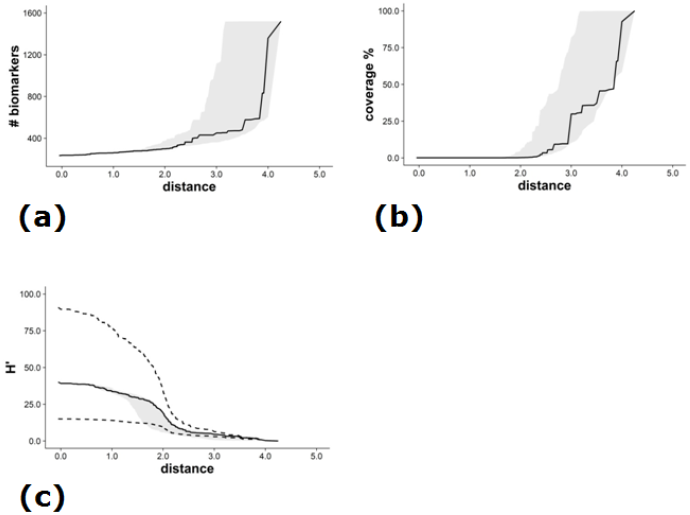
Biomarker characteristics across process aggregation levels. (a) Cumulative count of biomarkers involved in a single process or process aggregate in dependence of mean shortest path distance. (b) Fraction of median number of features represented by a biomarker and total number of features at a certain aggregation level (coverage, %). (c) Adjusted median Shannon entropy H’ (solid black line; first and third quartile given as broken black lines) for the set of 1,520 biomarkers in dependence of mean shortest path distance. For (a)-(c) the grey area is derived from controls resting on reference interactomes with reduced connectivity.

Among the 1,520 biomarkers 234 (15%) are assigned to a single process at L1 (the set of 3,860 processes), with a median number of seven assigned processes (the first and third quartile being three and 19). For biomarkers being member of more than one process at L1 consolidation into a unique, aggregate process develops at slow pace, at a distance of 2.1 seeing consolidation of 303 biomarkers (20%) in a single aggregate. The median number of process aggregates covered by a biomarker at distance 2.1 (aggregation level 1,177) is four (the first and third quartile being two and nine). The majority of biomarkers consolidates at a large distance of > 3.5, i.e. with the final ten process aggregate merge steps. 10% of biomarkers consolidate into a single aggregate only at the final merge event at level 1,340.

Biomarkers are intended as molecular process proxy, where process size expressed as number of molecular features involved is to be considered. The fraction of molecular features in processes holding a biomarker with respect to the total number of features involved at a specific aggregation level is shown in Fig. 2b. Up to a distance of 2.1 the fraction of features to be represented by a biomarker is about 0.2% of the total number of features involved at a given level, reaching a median value of 30% at a distance of 3.0 and a sharp increase with the final aggregation steps. Reference computation resting on interactomes with reduced connectivity indicates stability of curve characteristics for biomarker consolidation, feature coverage and entropy. For the latter a modified Shannon entropy expression H’ is derived. H’ aims at estimating the level-specific entropy of a biomarker with respect to the probability of populating microstates of a given process set (specific occupation of processes) when considering the process configuration at a certain level as the macrostate (combinatorics of process occupation) (Fig. 2c). The median dependence of H’ on distance exhibits a sigmoid behavior, with an inflection point at a distance of 2.1. As indicated by the quartile range individual biomarkers show distinct courses of entropy across aggregation levels, with selected examples shown in Fig. 3.

**Fig 3:**
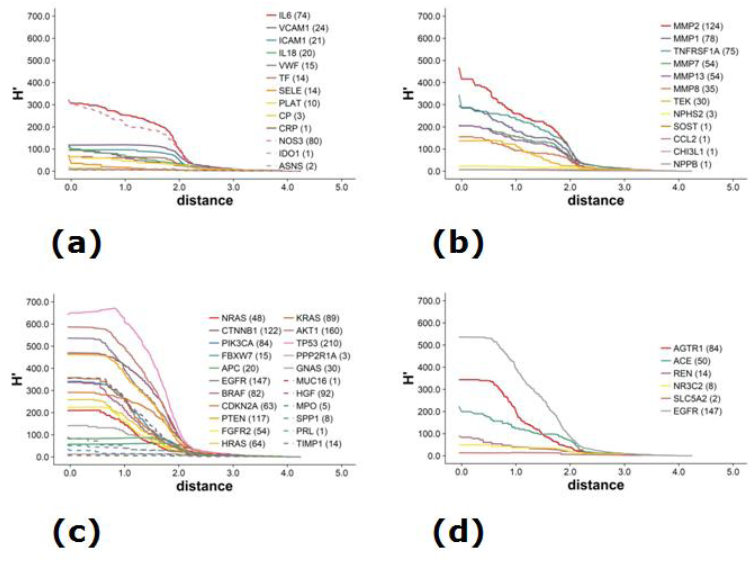
Information content of selected biomarkers. Adjusted Shannon entropy H’ for selected biomarker disease context including the number of processes embedding a biomarker at L1 provided next to the gene symbol: (a) Individual biomarkers in DN prognosis (solid lines) and drug response prediction (broken lines); (b) Biomarker panel in DN; (c) Members of a multi-analyte panel including amplicons (solid lines) and protein markers (broken lines) in cancer prognosis; (d) Selected markers serving as drug target.

Adjusted Shannon Entropy H’ is given for biomarkers and factors resembling drug targets in focus of two selected clinical indications, namely a diabetes co-morbidity (diabetic nephropathy, DN) and an example from oncology. Fig. 3a holds biomarkers with individual evidence for statistically significant associating with onset or progression of DN as reviewed in Hellemons et al. [17]. The biomarker listing includes eleven proteins (ten being covered in the biomarker core set of 1,520 features) and two metabolites. All but one marker exhibit a course of H’ in the quartile range of the total biomarker set considered (Fig. 2c). Outlier is IL6 (interleukin 6), being involved in 74 processes at aggregation level 1, consolidating into a single aggregate at aggregation step 1,339. Other extremes are CP (ceruloplasmin) involved in three, and CRP (C-reactive protein) involved in a single process at L1. For CRP an odds ratio (OR) of 1.06 is reported regarding onset and progression of DN, for CP the OR for onset of DN is 4.67 (at a sensitivity of 0.47 and a specificity of 0.84), the respective OR for IL6 is 1.72. The reported prognostic capacities include correction for conventional risk factors, and indicate insufficient power of these individual markers for assessing DN progression characteristics. DN is a complex disorder with involvement of multiple molecular processes from various compartments of the kidney, overall triggered by hyperglycemia and further impacted by micro- and macrovascular disease. As one strategy for respecting this complex background biomarker panels have been studied. Fig. 3b identifies biomarkers included in a classifier for predicting progression of DN [18]. The classification function was built using a least absolute shrinkage and selection operator (LASSO) for short-listing parameters which in combination add to explanation of variance of disease progression. For prognosis a combination of established risk factors (specifically baseline urinary albumin, systolic blood pressure and baseline estimated glomerular filtration rate (eGFR)) and given protein biomarkers provided a significant increase in R^2^ from 37.7% to 54.6%. Of the in total 13 biomarkers included in the classifier 12 are covered in the given core biomarker set. In contrast to the individual markers given in Fig. 3a all but three markers included in the panel see significantly higher start entropy, at a distance of about 2.0 meeting in entropy decay with the biomarker set discussed in Hellemons et al.. In the same context of progressive DN Pena et al. discuss a set of enzymes on the level of associated metabolites relevant in predicting response to angiotensin II receptor blockers (ARBs) in patients with diabetes mellitus [19]. Risk prediction of change in urinary albumin excretion (> 30% decrease) in response to ARB therapy provided an R^2^ of 0.5 when including selected protein biomarkers, in contrast to a R^2^ of 0.1 when using clinical parameters only. Entropies of respective protein markers again split into high (NOS3, with a start H’ of 324.05 and coverage of 80 processes at L1) and low entropy features (IDO1 and ASNS, with H’ and coverage at L1 of 6.30/1 and 8.62/2, respectively).

Evaluating a biomarker panel from a different clinical domain also identifies significant spread in H’ (Fig. 3c). Cohen et al. published a multi-analyte blood test for detection and localization of surgically resectable cancers [20]. The test rests on a classifier including assessment of abundance of circulating proteins and mutations identified in cell-free DNA. All 16 reported amplicons are covered in the given core biomarker set, for the protein set all but two analytes (the antigen on the tumor surface marker Sialyl-Lewis A, CA19-9; the carcinoembryonic antigen, CEA) are covered. The amplicons exhibit significantly higher entropies for distances up to 2.1 compared to the protein biomarker set. All but three amplicons (F-box/WD repeat-containing protein 7 FBXW7, Adenomatous-polyposis-coli protein APC, Serine/threonine-protein phosphatase 2A 65 kDa regulatory subunit A alpha isoform PPP2R1A) exceed the reference entropy distribution of the third quartile derived from the full biomarker set (Fig. 2c).

Selected biomarkers serving as drug targets are displayed in Fig. 3d. A first set focuses on targets embedded in the renin-angiotensin-aldosterone system (RAAS), with prominent drug classes used in DN, on a clinical observables level mainly intended for targeting vascular events [21]. High entropies are in particular identified for ACE (angiotensin converting enzyme) and AGTR1 (Angiotensin II Receptor Type 1), less pronounced entropy values are seen for REN (renin) and NR3C2 (aldosterone receptor). Associated drugs include ACEi (as Ramipril), ARBs (as Telmisartan), renin inhibitors (as Aliskiren) and mineralocorticoid receptor antagonists (as Spironolactone).

A different drug class focusing on a core mechanism driving DN as a diabetes associated disorder is SGLT2 inhibitors from the Gliflozin group. SGLT2 (SLC5A2, sodium/glucose cotransporter 2) is executing glucose re-uptake in the kidney proximal tubule segment. Inhibiting this mechanism targets hyperglycemia [22]. In terms of entropy, SLC5A2 is involved in only two processes at L1, consolidating into a single aggregate at L886.

A distinctly different behavior is identified for the epithelial growth factor receptor (EGFR) also being included in the multi-analyte panel tested in oncology indications [23,24]. EGFR as target, with major drugs including Cetuximab and Erlotinib, is embedded in 147 processes at L1, and shows one of the highest entropy courses of the given biomarker set.

## Discussion

Over the last decades most valuable, pharmacologically active compounds have become available, each holding extensive characterization of direct target interaction together with pharmacokinetics, dynamics and toxicology. However, subsequent attrition in clinical trials has remained in the range of about 50% in phase 1, 70% in phase 2 and 30% in phase 3 [25].

Many active compounds failing in a clinical trial (in particular at advanced trial stage) are not further followed, neither for evaluating alternative stratification within the same indication nor in repurposing, although alternative stratification may still bring benefit to unmet clinical needs. Exceptions include the Nrf2 pathway activator Bardoxolone methyl, initially considered in cancer indications before entering trials in DN. Worth to note is the repurposing rationale for Bardoxolone methyl. When tested in the oncology context an improvement of eGFR indicative for beneficial effect on renal function was seen. However, the BEACON clinical trial in type 2 diabetics with advanced chronic kidney disease did not reduce the risk for end stage renal disease (ESRD) or death from cardiovascular events. The latter endpoint even saw an increase in the treatment group leading to termination of the trial [26]. Still, post-hoc analysis concluded on preservation of kidney function as determined on the level of eGFR [27], triggering further trials in kidney disease. Positive phase 2 clinical trial results were reported from the CARDINAL study utilizing Bardoxolone for improving eGFR in Alport Syndrome (a genetic condition also affecting kidney function), i.e. using the same clinical surrogate and drug mechanism of action for addressing a fairly different disease etiology and pathophysiology. Trial results see a debate about competing surrogate markers, as next to an increase in eGFR an increase in the urinary albumin-to-creatinine ratio (UACR) was identified, being considered as indicator for kidney disease [28]. Eventually, inclusion of further biomarkers aside higher order clinical parameters may improve information content for optimizing stratification.

For approved drugs comprehensive data on their benefit in clinical reality have become available. For example, the ACCORD study evaluated the effect of targeting a systolic blood pressure of less than 120 mm Hg in contrast to less than 140 mm Hg in patients with type 2 diabetes, concluding on no reduction of a composite outcome (fatal or nonfatal major cardiovascular events) [29]. On the other hand, multi-factorial intervention (addressing glucose and lipid levels together with blood pressure) identified significant reduction of cardiovascular events in a cohort of type 2 diabetes patients with microalbuminuria [30].

Davis et al. captured cancer drugs approved by the European Medicines Agency (EMA) in 2009-2013 (in reference to a U.S. FDA cancer drug evaluation for approvals covering 2008-2012) [31]. The authors conclude that after a minimum of 3.3 years post market entry evidence on benefit or improvement of quality of life compared to given treatment options is limited for a substantial fraction of such drugs. Among 68 cancer drug indications eleven were considered exhibiting a clinically meaningful benefit. The authors further note limited additional information being generated post approval to guide clinicians. One of the central challenges seen is use of surrogate parameters with limited evidence for linking with endpoints.

The situation is further complicated when asking for patient-specific benefit aside identified significance on a patient cohort level. A trial in DN utilizing the ARB Irbesartan demonstrated a clinically meaningful reduction of a composite endpoint (doubling of serum creatinine, development of end-stage renal disease or death from any cause) by 20% in a follow-up period of 2.6 years [32]. However, at present no tools are available for identifying the group of responders at initiation of treatment, and 50% of patients in the treatment group reached the primary endpoint after 54 months.

Aside given evidence in relating clinical observables with disease progression and outcome, and beneficial effect of drugs on a cohort level, statements on personalized drug response and precision in drug use apparently fall short. Still, clinical phenotyping remains the main route in defining clinical trial inclusion criteria. For instance, at present 28 clinical trials are identified for DN in the clinicaltrials.gov registry of clinical trials (interventional, recruiting or active, phase 2 and 3). 27 trials rest on established clinical phenotyping inclusion criteria. Just one trial involves a molecular profiling approach (PRIORITY trial) [33]. This design uses clinical phenotyping (established type 2 diabetes mellitus, eGFR >45 ml/min/1.73m2, UACR < 30 mg/g) for defining a patient population susceptible for developing DN. Next, a proteomics-based classifier for further stratification with respect to disease onset and progression is applied. If the resulting stratum resembles a homogenous phenotype on the pathology level (next to progression characteristics) specifically responsive to Spironolactone mechanism of action (primary endpoint being confirmed microalbuminuria defined as UACR >30 mg/g) is to be demonstrated.

Clinical phenotyping for diagnosis and prognosis, together with parameters used for assessing drug response, resemble higher order observables. This parameter class captures overall characteristics of pathophysiologies originating from a molecular architecture approximated as process-of-processes configuration. Such parameters, individually or in combination, are used for defining a specific phenotype instance. However, with this approach multiple genotypes (defining individual patients or groups of patients) are assigned to a certain phenotype. With involvement of different genotype instances the molecular organizational representation is heterogeneous. As the molecular architecture is the context being affected by a drug’s mechanism of action, valid mapping to drug response is per conceptual design limited.

eGFR serves as example, being a purely functional indicator for kidney filtration capacity (clearance of plasma creatinine, in an empirical expression further including age, race, gender), not informing on specific kidney pathology. For improving phenotype assignment eGFR is combined with information on disease etiology (as diabetes, or genetic testing combined with histopathology from kidney biopsies in Alport Syndrome), eventually combined with further parameters for improving prognosis. Still, such parameter combinations may in some sense define a homogeneous phenotype, but not a representative hyperstructure, in consequence leading to variance in drug response. A frequently pursued analysis strategy for yielding improved parameter combinations even impedes improved resolution of molecular architecture, namely evaluating if parameters add prognostic or predictive power on top of established higher order readout. In such approach molecular surrogates of established clinical parameters are eliminated, in consequence limiting insight into respective molecular process configuration. As example, a molecular marker adding on top of eGFR may improve prognosis and also add information on a specific molecular context, but leaves all variance explained by eGFR unresolved on a molecular mechanism level. Such approach does not contribute to matching drug mechanism of action.

This context introduces a study design issue: A cohort defined on the level of higher order readout apparently embeds more than one hyperstructure instance. Supervised analysis per design leads to probabilistic expressions on the level of individual patients. In contrast, unsupervised analysis may guide in separation of embedded hyperstructure instances. Such approach was recently pursued in type 2 diabetes mellitus [34]. Utilizing six clinical observables allowed dissection of five distinct phenotypes with respect to developing diabetes complications, with one cluster specifically prone to developing DN. Further, patient cluster-specific genetic associations could be retrieved. The authors conclude that such sub-stratification may enable first steps toward precision medicine in diabetes. According to the hyperstructure concept subsequent steps may involve identification of biomarkers characterizing molecular process configuration specifics of subgroups being identified in unsupervised analysis for allowing tailored matching of drug mechanism of action.

Considering a hyperstructure as valid proxy of process organization resembling pathologies the question arises on what specific organizational level optimal information content of a parameter is to be expected. For instance, given data on DN show limited information gain on the context-free level of genetic traits (in the hyperstructure notion being at level 0). Effect size seen with genetic variants explains only a limited range of phenotype variability. Still, the genetic component may well present with more subtle readout on the level of processes, specifically when meeting with parameters as hyperglycemia. Following this argument, biomarkers resembling proxies of molecular process configuration may hold improved information content compared to genetic traits.

The given approach for deriving a human pathology hyperstructure as set of process configurations involves a number of substantial approximations. Annotation of pathologies is limited to available and accessible information on molecular features associated with disease terms according to the NCBI MeSH vocabulary, presumably being far from complete. The second major approximation is delineation of molecular processes on the level of given interaction network characteristics, with shortest paths as criterion for process formation and interaction. Thirdly, hyperstructures resemble a dynamical context to be derived bottom-up, whereas the given pathology hyperstructure confers to a static approximation.

Aside these limitations the hyperstructure derived from a molecular process set covering 1,008 phenotypes shows distinct characteristics. With 3,860 processes as start condition 238 k-cores aggregate at distance zero, i.e. more than 20% of the phenotypes being covered hold shared molecular aspects. Significant redundancy is also displayed on the level of molecular features, where the unique set of about 5,000 protein coding genes expands by a factor of about seven at the first level of process configuration. Median number of features involved in a process at level 1 is five (first quartile of three, third quartile of ten). The number of processes sees a steep decline along aggregation up to a mean shortest path distance of 2.1 (aggregation level 1,177), where 290 processes remain. With a reduction of processes by 90% feature redundancy is still at a factor of about three compared to the non-redundant feature set. Median number of features embedded in a process aggregate at this level is 23 (first quartile of nine, third quartile of 60 features). The nominally small values of mean shortest paths are mainly driven by feature overlaps across processes, according to process distance computation contributing with a path length of zero. Reporting biases may to some extent contribute to feature redundancy, however, with a set of about 1,000 leaf terms distributed across all major branches of the MeSH disease ontology this bias may be limited. Further aggregation of the 290 processes identified at aggregation level 1,177 takes place in 163 merge steps, with the final aggregate holding a shortest path of about 4.2. This path length is also driven by hubs in the scale free topology of the reference network, in the context of disease pathologies defining the lower bound of path length. Organ and tissue-specific absence of node activity will increase effective path length, but eventually not seeing a significant change due to essentiality of hub nodes.

Redundancy is equivalently reflected for features holding biomarker annotation. First to note is the large number of protein coding genes holding annotation in the diagnostic or prognostic context. For about one third of features involved in the process set at least one co-notation in the set of 1,008 disease terms is given, however, with large variation. 50% of disease terms hold no biomarker annotation, whereas other individual terms specifically in the oncology domain see large numbers of assigned biomarkers. These trends are also reflected on the level of individual features, with e.g. TP53 or AKT1 being assigned as biomarker to a substantial number of disease terms.

On a process level redundancy further expands, as a feature holding annotation as biomarker may be considered relevant in certain pathologies and occur in process sets of other pathologies aside actual biomarker annotation in such diseases. With a median of seven processes at level 1 and four aggregate processes at level 1,177 multiple involvement of biomarkers in pathology processes across the hyperstructure is evident. This status is reflected in modified Shannon entropy values H’ for the biomarker set via respecting distribution of a biomarker across level-specific process sets, additionally taking the size of processes on the level of embedded features into account. For the given biomarker set of 1,520 features median H’ values remain largely constant up to an inflection point at distance 2.1. Apparently, median entropy and respective information content of the biomarker set does not substantially change for a wide range of aggregation levels.

Individual biomarkers show distinct behavior in entropy at process start configuration and also in respective decay. A first set assigned to DN covers individual biomarkers, each exhibiting statistically significant association with onset or progression of the disease. Nine out of ten markers display moderate start entropy, with ICAM1, VCAM1 and TF seeing minor change in entropy up to aggregation level 1,177. In contrast IL18, VWF, PLAT and SELE see entropy decline and respective consolidation already at earlier stages of process aggregation. A different behavior is observed for IL6, with broad distribution across processes at the start configuration and final consolidation in an aggregate process close to maximum distance. IL18, VWF, PLAT and SELE are reported as associated with onset and progression of DN, however, with odds ratios or correlation coefficients being in a range lacking clinical relevance.

A different entropy level is seen for a second set of biomarkers tested for progression of DN, with on average substantially higher entropies. Biomarkers reported as significantly associated with eGFR decline in univariate analysis and showing specifically high entropy include TNFRSF1A, MMP2 and MMP7, complemented by the low entropy markers CHI3L1 and NPHS2. In multivariate analysis joined with clinical phenotyping parameters top ranked markers with respect to explaining variance in eGFR are MMP2, TEK, MMP7 and MMP8, all being high entropy markers on the basis of the given hyperstructure.

A distinct split in low- and high entropy markers is also seen for the oncology panel covering amplicons and protein markers, the latter showing particularly low entropy with exception of HGF. In contrast, amplicons covering generic effectors from cell cycle and differentiation include the maximum entropy marker TP53.

Spread in entropy is also seen for markers utilized as drug targets. For addressing RAAS, AGTR1 as well as ACE display high entropy, in contrast to the aldosterone receptor NR3C2. ACE inhibitors and angiotensin II receptor blockers are among the most frequently prescribed medications in the cardiovascular context including DN, apparently addressing targets displaying high entropy across all aggregation states. Very much in contrast is entropy of a novel approach in tackling diabetes complications via targeting SLC5A2, a highly specific target in the renal proximal tubular compartment. An equivalent status holds NPHS2 (Podocin) included in the classifier for assessing DN progression. Low entropy of this feature confers to highly specific expression and function in kidney podocytes.

## Conclusions

Clinical laboratory routine diagnostics offers > 2,000 parameters, including a limited number of individual protein markers. Examples include cardiac markers as proBNP (NPPB, low entropy) and Troponin T (TNNT2, medium entropy), or features also involved in the panels discussed above (as IL6, CEA, CREA, ACE, REN, MUC, ICAM, VCAM, CRP, VWF). Lack of validation joined with study design issues limits further inclusion of biomarker candidates proposed from discovery studies, as clinically relevant sensitivity and specificity is not demonstrated, even more so when it comes to drug response prediction. In further consequence development of companion diagnostic assays for adding precision in drug use stalls.

With biomarker analysis on the pathology hyperstructure the following hypothesis is derived: For adding precision in predicting drug response a biomarker panel grounding in disease-associated processes needs to include high and low entropy markers. High entropy markers serve as molecular surrogate for systemic consequence, and low entropy markers add specifics of target pathology. Combining both biomarker classes promises capturing essential characteristics of molecular process configuration, and in consequence improved assessment of drug effect.

A biomarker solely embedded in a single process at level 1 of the hyperstructure is specific for such process, but may offer limited information about process context. It appears fair to assume that clinical phenotype parameters only come into existence at higher order process configurations. In analogy, biomarkers with higher entropy resemble proxies of higher order readout, at some level meeting with information content seen in clinical phenotyping.

The commonly pursued strategy in biomarker screening focuses on identification of most specific processes for a pathology. Given analysis results indicate that such markers need to be combined with features displaying increased entropy for finally adding in precision when it comes to drug response prediction.

Equivalent reasoning may be applicable for drug targets, with the high entropy features ACE and AGTR1 as examples. With clinically relevant benefit of ACEi and ARBs, the number needed to treat is identified as suboptimal, a fact eventually grounded in limited information content of these targets. According to the given hypothesis the higher order function readout of these features needs to be combined with low entropy markers reflecting specific consequences of elevated blood pressure, be it in kidney-, micro- or macrovascular pathology and corresponding clinical events. With the inflection point in process aggregation and biomarker entropies a relevant resolution of process aggregates may be given where information content on a macrostate level meets with optimal resolution for patient-specific microstate configurations. Eventually such entropy considerations support design of biomarker panels for improved phenotyping and in consequence improve tools for adding precision in drug selection and response prediction.

## Methods

### Formal considerations

A disease phenotype manifests as clinical observables, and grounds in molecular architecture of disease pathology. Whereas for most phenotypes clinical characterization is far from exhaustive, available clinical observables serve diagnosis of a disease together with prognosis with respect to clinical events and outcome, in consequence defining treatment regimens. Evidence for linking observables and outcome mostly grounds in empirical, observational studies. Clinical trials on the other hand resemble the gold standard for providing evidence on beneficial effect of a treatment on outcome (or surrogates of outcome) when using certain observables as inclusion criteria. A relation of observable and outcome may be expressed as marker or factor, with the latter exposing a causal relation of pathology and phenotype specifics. As consequence, factors are relevant for prognosis of disease progression, may serve as surrogate of outcome, and identify relevant drug target molecular context. Surrogates are considered essential for improving treatment modalities specifically for chronic diseases. Such parameters enable executable clinical trial designs in case of extended time-to-event spans, and may allow intervention at earlier stages of disease pathology eventually seeing improved likelihood of beneficial effect. With respect to treatment response, factors further serve as monitoring element and provide predictive measure. Specifically in the latter case the difference of marker and factor becomes evident, where a marker changes e.g. in abundance level with treatment, but this variation does not associate with outcome.

In most clinical conditions treatment effect is linked with observables and only in case of factors links with outcome. However, when it comes to treatment and its molecular mechanism of action the quest is in establishing a relation of phenotype and molecular pathology, the latter being the molecular context of treatment mechanism of action. In formal terms, some expression is to be established allowing a mapping of a molecular status (a genotype resembling certain pathology) and a phenotype, eventually mitigated by observables according to the scheme given in Fig. 4a.

**Fig 4:**
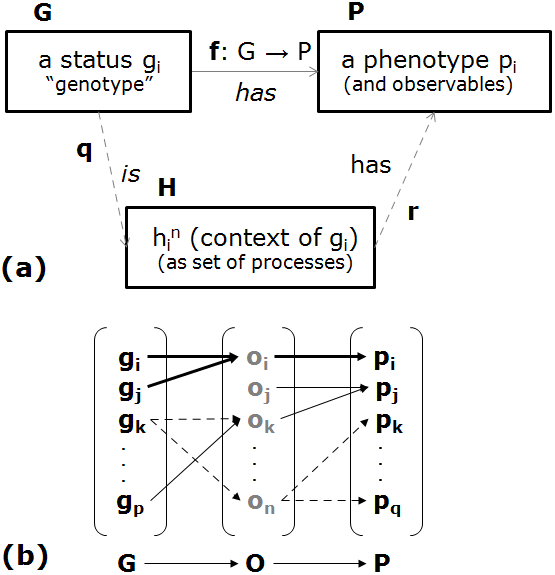
Relation of genotype, phenotype and observables. (a) Mapping of genotype set **G** and phenotype set **P** (with selected instances **g_i_**, **p_i_**) and an alternative mapping via hyperstructures from the set **H** (and an instance of order **n**, **h_i_^n^**). **f**, **q**, **r** are functions. (b) Valid (solid line) and non-valid (dotted line) mappings of instances in **G**, **O** and **P**.

**G** is the set comprising genotype instances **g_i_**, …, **g_p_**. When it comes to instances resembling a certain clinical presentation the true genotype space **G** is at present not defined. The same holds for the set of distinct phenotypes **p_i, …_ p_q_**, as available observables lack necessary resolution. With no exhaustive enumeration of **G** and **P** a closed form delineation of a functional relationship **f** is not amenable. Therefore, **f** is in practice replaced by statistical expressions derived on a patient cohort level, involving characteristics from **G** and observables from clinical phenotyping e.g. in regression functions, classifiers, but also single parameter cutoff values (as stage classification in kidney diseases according to eGFR). Formally, **f** may assign multiple instances of **G** to a single **p_i_**, but no single **g_i_** shall map to more than one instance of **P**, otherwise **f** is not a valid mapping. When using statistical approaches this condition is not satisfied. The consequence is a probabilistic assignment of phenotype instances, per construction introducing variance.

Whereas there exists the mapping of each **g_i_** to some **p_i_** (as every patient holds a defined genotype presenting as defined phenotype), procedures utilize observables from clinical phenotyping as intermediate. According to Fig. 4b both, **g_i_** and **g_j_** map to a single observable **o_i_**, whereas the mapping of **g_k_** to both, **o_k_** and **o_n_** is not valid; the same rationale holds for the non-valid mapping of **o_n_** to **p_k_** and **p_q_**, whereas **p_j_** is validly defined via **o_j_** and **o_k_**. Hence, a valid element of **f** for **X** → **P** is {**g_i_**, **g_j_**} → **o_i_** →**p_i_**. In such notion, from 1 to **m** realized genotype instances display a distinct observable being sufficient for mapping to the respective phenotype instance. Important to note, such observable may be a single parameter or a parameter combination (a panel). Such observable **o_i_** is deemed to provide diagnostic and prognostic information, together with educating on relevant mechanism of action of treatment with respect to the involved genotype instances. Precision medicine approaches are centered on such observables, in stratification factoring in 1-m genotype instances. With the limit m=1 in personalization, implementation in drug development and clinical practice appears impractical. The ideal observable aims at identifying a maximum number **m** of genotypes for which a mapping to a single **p_i_** is still valid.

For pursuing precision strategies on the basis of stratification the characteristics of such class of observables needs attention. Each genotype instance itself is composed of a set of components in interaction, in biological terms resembling the coding regions of a given genome. Each such element holds properties as certain biological function reflected in domain structure, becoming apparent on the respective RNA or protein effector level. We denote these components as being of order zero, each exhibiting observables of order zero, **O^0^**. Examples for experimentally amenable observables include variation in functional properties (e.g. caused by genomic alterations as single nucleotide polymorphism). Another property of such components also reflected in sequence and respective structural domain composition is interaction with other components, by this defining component context. Including context on the level of zero order component ensembles leads to a frequently used system representation in the form of interaction networks.

Components in such context apparently exhibit organizational structure in the form of subgraphs termed molecular processes, and via formation of structure new system properties and associated observables of first order, **O^1^**, are displayed. Such first order structures form further process context, including as examples apoptosis, senescence or proliferation, resembling the consequence of organizing processes in pathways. Such second order structures assemble into yet higher order organizational structure and concomitant observables, finally exhibiting clinical phenotype parameters, **O^n^**, with e.g. eGFR or blood pressure as representative instance. While these examples cover a bottom-up construction principle also a top-down effect needs to be taken into consideration. For example, among other effects blood pressure contributes via induction of endothelial cell stress to provision of lower order observables as arterial sclerosis, in turn triggering cardiovascular events as high order observable. Such system construction involving up- and downward causalities resembles dynamical hierarchies holding configurations of process sets as component-in-context phenomenon, and forms hyperstructures **h_i_** of order **n** [35,16] (Fig. 5). In formal terms a hyperstructure **h_i_** of order **n** under a dynamics **R** capturing interaction of components (**Int**) is given as:

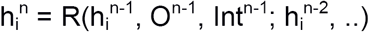

**Fig 5:**
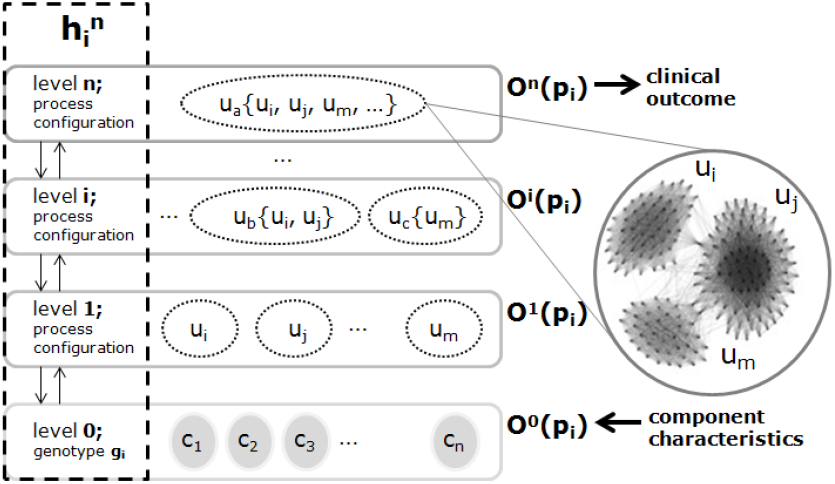
Schematic representation of a molecular process hyperstructure **h_i_ ^n^** and observable classes of a phenotype instance **p_i_**. Starting with an instance of **G** (**g_i_**, holding components c_1_, …, c_n_, and respective observable class in **O^0^**) molecular processes u_i_, … u_m_ aggregate into process configurations involving up- and downward causalities across all complexification levels up to **n**. Process configuration (schematically indicated as network model of three processes) finally determines high order observables, ultimately involving the level of clinical outcome.

Accordingly, for mapping genotype instances to a phenotype instance we seek an observable serving as proxy for common aspects of a hyperstructure **h_i_**(**p_i_**). The ideal observable captures specifics of a hyperstructure, with the assumption that the dimension of **H** is significantly smaller than the dimension of **G**, and equal to the dimension of **P**. Stratification seeks for a hyperstructure **h_i_** capturing a subset of genotypes exhibiting the same phenotype **p_i_**.

Three distinct classes of observables appear amenable in practice, being either in **O^n^**, in **O^0^**, or in between these limits. The first class covers consequence of forming organizational structure, as such not being displayed as property of zero order objects. Such observables involve most of present clinical phenotyping. A second aspect from the same class of observables covers metabolites, whose abundance on a molecular level mirrors the state of molecular process configuration (as blood glucose or lipid levels), although themselves not being components of a genotype. The second class of observables, given in **O^0^**, covers molecular peculiarities of genotype instances as e.g. genetic traits. The third class covers dynamical markers responsive to molecular context, for instance abundance of protein biomarkers.

With cumulative interaction of processes forming a hyperstructure the definition of biomarkers comes in handy, namely being a characteristic that is objectively measured and evaluated as an indicator of normal biological processes, pathogenic processes, or pharmacologic responses to a therapeutic intervention. In the formalism of dynamical hierarchies of processes the following assumptions appear valid: i) each observable in **O^n^** holds a proxy on the level of **O^>0^** (with the exception of certain genetic disorders even holding a direct proxy at **O^0^**) being either a property of an individual molecular feature or a combination of properties of more than one feature, giving rise to marker panels; ii) not all specifics of a hyperstructure provide an observable in **O^n>0^** being effectively amenable for quantitative measurement; iii) most observables in **O^n^** being used in clinical practice hamper a unique assignment to a phenotype and respective hyperstructure instance resembling the root cause for lacking precision in treatment. In contrast, molecular biomarker properties in **O^>0^** may allow a valid mapping if resembling a proxy for a molecular process configuration. The latter assumption provides a direct link with drug molecular mechanism of action. Drug effect aims at re-configuration of a given pathology hyperstructure for altering the result of the phenotype mapping, e.g. rendering a progressive phenotype into a stable presentation.

Key element is search for observables in **O^>0^** which are sensitive and specific for molecular process configuration, instead of solely providing information on individual molecular process status. Such biomarkers reflect molecular context of a hyperstructure configuration, hence serve as proxy for higher order readout. Replacing **O^n^** with molecular context proxies in **O^>0^** promises improved representation of **h_i_**, and in consequence improved precision for mapping to **p_i_**.

### Workflow for establishing molecular process hyperstructures

According to the definition of a hyperstructure cumulative aggregation of pathology-associated molecular processes is to be derived, followed by evaluation of biomarker information content allowing to monitor higher order readout established by process configuration.

As depicted in Fig. 6 start elements involve a reference interactome and a set of disease terms. The interactome holds protein coding gene-centric nodes, and interactions combine evidence from experimental sources and computational inference. The set of disease terms is derived from an established controlled vocabulary, for each term retrieving molecular features reported as relevant in disease pathology. Combining network and molecular disease annotation provides network models (subgraphs) of pathologies, on such basis deriving molecular processes as core pathology components. Aggregating the processes in a static representation of a hyperstructure allows evaluation of information content of disease term-associated biomarker candidates.

**Fig 6:**
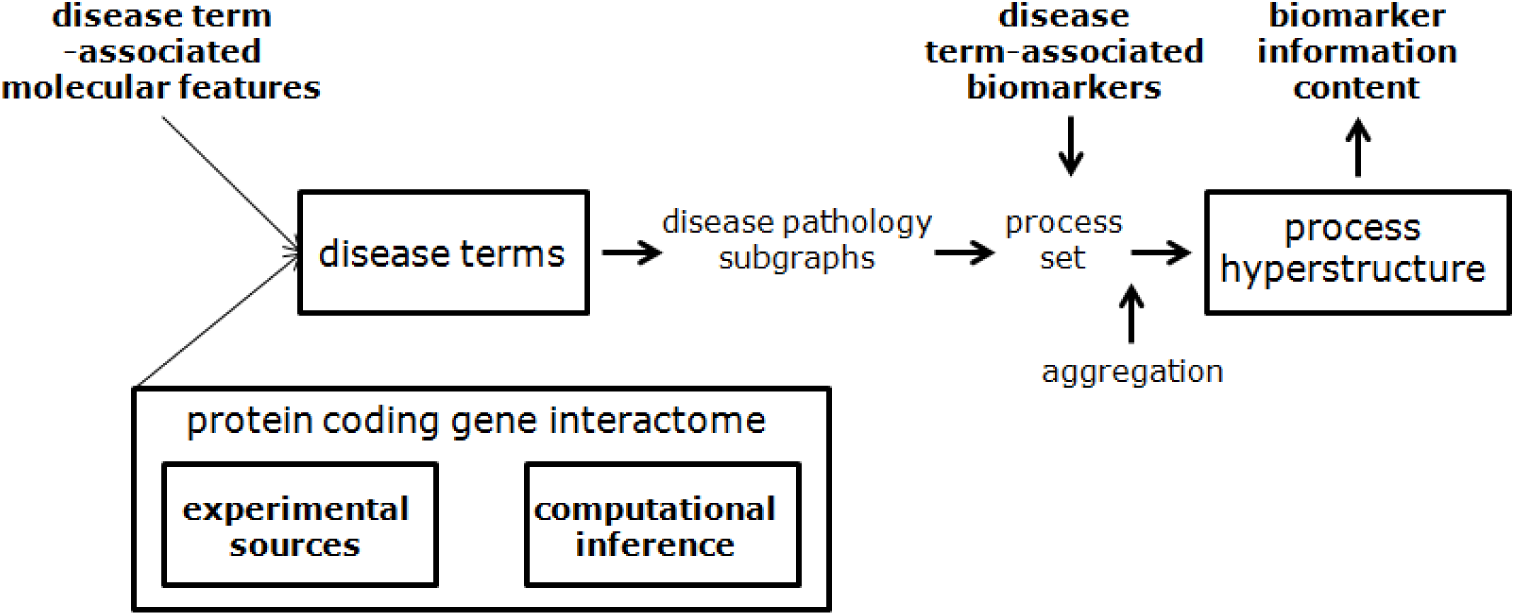
Workflow scheme for establishing a static approximation of a molecular process hyperstructure capturing human pathologies for deriving biomarker information content.

#### Interaction network, pathology process models and biomarkers

A human protein coding gene interaction network serves as basis for pathology process models. The reference network integrates experimentally derived interactomics data and computationally inferred interactions, covering 17,671 human protein coding genes according to Ensembl [36] as nodes and 705,283 interactions as edges [16]. For human protein coding genes as defined on the level of Gene Symbols interaction data are retrieved from BioGRID/362,951 interactions [37], IntAct/264,836 interactions [38] and Reactome/212,513 interactions [39]. The interactions cover data points from experimental procedures (interactomics assays in various formats) together with expert curation. Consolidating interaction sources results in 511,184 unique interactions (from the total number of 840,300 interactions).

For addressing the false negative rate of the given human protein coding gene interactome this reference interaction set is expanded via computational interaction inference as detailed in Fechete et al. [40]. Briefly, annotation from at least one of the data sources InterPro [41], GO Function and Process [42], Reactome and Panther pathways [43] was consolidated, using the longest protein isoform as canonical sequence of the respective protein coding gene. For each pair of genes a score was derived for an interaction of type “procedural” (composed of GO process together with Panther and Reactome pathway information) and of type “functional” utilizing annotation information from GO function together with protein domain information as retrieved from InterPro. The computed scores, normalized in the interval [0,1], correspond to a probability of establishing an interaction. Accuracy of the approach was tested against experimentally available data from an independent source (KEGG, [44]), with maximum precision reached at a score cutoff value of 0.74. This procedure (implemented as Python routine) provided additional 194,099 inferred interactions.

Computation of molecular process models approximating disease pathologies integrates information on molecular features associated with pathologies and the reference protein coding gene interaction network as detailed in [16, 45]. The annotation procedure utilizes NCBI Medline, offering citations with annotation in Medical Subject Headings (MeSH) disease terms [46]. A further source provided by NCBI is gene-to-PubMed [47], listing relevant protein coding genes for a given publication. Combining both allows consolidation of molecular features reported in the context of individual disease terms. For each disease term defined in MeSH associated PubMed identifiers (from a total set of about 27 million publications) were retrieved. For each such publication a lookup in gene-to-PubMed identified protein coding genes as applicable, subsequently being assigned to a disease term in focus. This routine was executed for all 3,066 leaf disease terms listed in MeSH.

With disease term-associated molecular features as well as nodes of the reference interactome being on the level of protein coding genes defines disease term-specific subgraphs. Each such subgraph restricts the node set of the interactome to the protein coding gene set retrieved for a disease term together with interactions among such nodes according to the reference interactome. Forwarding a subgraph to the cluster identification algorithm MCODE [48] allows extraction of densely connected regions (k-cores resembling subgraphs of minimum degree k), leading to a topology-based definition of molecular processes. A molecular pathology model of a given disease term is then considered as the set of k-cores (processes) retrieved from the disease term-specific subgraph.

Finally, protein coding genes holding annotation as biomarker are derived from NCBI MeSH utilizing the descriptor “Biological Markers” in the specific categories “diagnosis”, “prognosis” in conjunction with a disease term as defined in MeSH [16,49].

#### Molecular process aggregation and network measures

For a given disease term zero to **m** processes (k-cores) are derived, resulting in a set of **n** processes across all disease terms holding at least one process. A distance between all pairs of processes is calculated on the basis of the reference interaction network. For each process pair irrespective of specific disease term assignment the average shortest path length from all nodes of the first process to all nodes of the second process is determined. In case of features being present in both processes the shortest path equals zero. The procedure provides a pair-wise process distance matrix, and serves as input for complete linkage hierarchical clustering (see Additional file 1). The Scientific Computing Tools for Python (version 1.0.0) is used for clustering, downstream analysis and visualization of the dendrogram [50]. For evaluating stability of aggregation characteristics 100 reference runs are executed at reduced network connectivity. In each such run 30% of given interactions (the fraction of inferred interactions added to interactions holding experimental evidence) of the reference network are randomly removed, followed by computation of the hierarchical clustering and respective process aggregation and biomarker characteristics. As maximum process distances increase with removal of interactions all computed characteristics for control networks are normalized to the maximum path length of 4.2 seen with the reference network.

As biomarker entropy measure the Shannon entropy serves as basis [51], determined as *nat* (natural unit of information using the Euler number *e* as base of the natural logarithm). A modified Shannon entropy H’ is defined as a proxy of the information carried by the event of observing a specific set of process aggregates with a specific assignment of a biomarker together with all other process aggregates defining the macrostate, further taking into consideration the size of aggregates in terms of number of molecular features included.

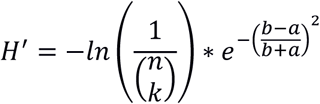

By using the binomial coefficient the number of possible configurations is calculated for reflecting the probability of observing with a single feature a specific configuration of processes of size k (number of processes containing the biomarker) out of all processes (n) seen at a certain process aggregation level. Equal probability of a feature to be contained in a process is assumed for the given setting. Each disease term is considered as independent with respect to its assigned process set, as reflected by high redundancy of features across the entire set of 3,860 processes. This situation is in contrast to a process definition originating from molecular pathways, where conditional probabilities appear preferable.

With the exponential term entropy is adjusted for deviating numbers of molecular features contained in aggregates. Conceptually, a biomarker serves as proxy for the status of a process independent of the number of features involved. However, information content in practice will see variance in truly capturing all information flow realized in processes with increasing number of embedded features. As approximation we use mean number of features of process aggregates not holding the biomarker, given by the parameter **a**, while mean number of features of process aggregates holding the biomarker is given by the parameter **b**.

## List of abbreviations

ACEi: Angiotensin Converting Enzyme inhibitor
ARB: Angiotensin II Receptor Blocker
DN: Diabetic Nephropathy
eGFR: estimated Glomerular Filtration Rate
EMA: European Medicines Agency
ESRD: End Stage Renal Disease
MeSH: Medical Subject Headings
NCBI: National Center for Biotechnology Information
NME: New Molecular Entity
NNT: Number Needed to Treat
OR: Odds Ratio
RAAS: Renin Angiotensin Aldosterone System
UACR: Urinary Albumin to Creatinine Ratio
U.S. FDA: United States Food and Drug Administration

## Availability of data and materials

The datasets analyzed during the current study are available at:

BIOGRID repository, https://downloads.thebiogrid.org/BioGRID ^37^

IntAct repository, https://www.ebi.ac.uk/intact/downloads ^38^

Reactome repository, https://reactome.org/download/current/interactors/reactome.homo_sapiens.interactions.psi-mitab.txt ^39^

Gene Ontology repository, http://geneontology.org/page/download-ontology ^42^

Panther Classification System repository, ftp://ftp.pantherdb.org/pathway/current_release/ ^43^

Medical Subject Headings repository, https://www.nlm.nih.gov/mesh/filelist.html ^46^

gene2pubmed repository, ftp://ftp.ncbi.nlm.nih.gov/gene/DATA/gene2pubmed.gz ^47^

## Authors’ contributions

BM designed the study. AL executed biomarker analysis. AH derived network models. All authors contributed to writing and approved the final manuscript.

